# Rapid microbial interaction network inference in microfluidic droplets

**DOI:** 10.1101/521823

**Authors:** Ryan H. Hsu, Ryan L. Clark, Jin Wen Tan, Philip A. Romero, Ophelia S. Venturelli

**Affiliations:** Department of Biochemistry, University of Wisconsin-Madison, Madison, WI 53706; Department of Bacteriology, University of Wisconsin-Madison, Madison, WI 53706; Department of Chemical & Biological Engineering, University of Wisconsin-Madison, Madison, WI 53706

## Abstract

Microbial interactions are major drivers of microbial community dynamics and functions. However, microbial interactions are challenging to decipher due to limitations in parallel culturing of sub-communities across many environments and accurate absolute abundance quantification of constituent members of the consortium. To this end, we developed <u>M</u>icrobial Interaction <u>N</u>etwork Inference in microdroplets (MINI-Drop), a high-throughput method to rapidly infer microbial interactions in microbial consortia in microfluidic droplets. Fluorescence microscopy coupled to automated computational droplet and cell detection was used to rapidly determine the absolute abundance of each strain in hundreds to thousands of droplets per experiment. We show that MINI-Drop can accurately infer pairwise as well as higher-order interactions using a microbial interaction toolbox of defined microbial interactions mediated by distinct molecular mechanisms. MINI-Drop was used to investigate how the molecular composition of the environment alters the interaction network of a three-member consortium. To provide insight into the variation in community states across droplets, we developed a probabilistic model of cell growth modified by microbial interactions. In sum, we demonstrate a robust and generalizable method to probe cellular interaction networks by random encapsulation of sub-communities into microfluidic droplets.

## INTRODUCTION

Microbial communities have a tremendous impact on diverse environments ranging from the human body to the plant rhizosphere (Berendsen et al., 2012; Clemente et al., 2012). Microbe-microbe and environment-microbe interactions are major determinants of microbial communities and microbiomes (Cao et al., 2018; Venturelli et al., 2016). Deciphering interaction networks in high-dimensional microbial communities is challenging due to the need to rapidly and accurately determine the absolute abundance of each community member across many sub-communities and environments (Cao et al., 2017; Harcombe et al., 2016).

The population sizes of microbial consortia can range from less than ten cells in mixed species biofilm aggregates to 10^11^ cells mL^-1^ in the human colon (Connell et al., 2014; Sender et al., 2016; Stoodley et al., 2001). Cellular growth history, the temporal order of strain colonization or the initial phase of microbial competition can impact community assembly (von Bronk et al., 2017; Kong et al., 2018; Vega and Gore, 2017; Venturelli et al., 2018; Zhou et al., 2013). Our understanding of microbial consortia in small populations is limited due to technical challenges in the manipulation and analysis of small populations of cells (Connell et al., 2014). Therefore, high-throughput methods that can rapidly resolve microbial interaction networks across different initial community states, population sizes and environments would enable a better understanding of the key parameters shaping the structure and function of microbial communities and how to harness these systems for diverse biotechnological applications.

Microbial interaction network inference requires accurate measurements of the absolute abundance of each member of the community (Fisher and Mehta, 2014). Recent experimental efforts have used models trained on measurements of 1-3 member communities to predict community composition or function of up to 12 members to varying degrees of accuracy (Friedman et al., 2017; Guo and Boedicker, 2016; Kong et al., 2018; Mounier et al., 2008; Venturelli et al., 2018). Absolute abundance quantification of each member of a microbial community has ranged from low-throughput selective plating to count colony forming units (tens of samples per experiment) (Mounier et al., 2008) to optical density multiplied by relative abundance based on next-generation sequencing of samples generated through robotic high-throughput culturing (hundreds of samples per experiment) (Venturelli et al., 2018).

Encapsulation of microbial communities into microdroplets has been used to study ecological and evolutionary processes of microbial communities (Bachmann et al., 2013; Park et al., 2011). Water-in-oil droplets can be generated at kilohertz (kHz) rates using microfluidics, wherein cells from a mixed culture are randomly encapsulated into droplets yielding diverse sub-communities that can be studied in parallel (millions of samples per experiment). Each droplet is a miniaturized compartment that can be used to study interactions between community members in small populations. Microfluidic technologies enable the generation of well-controlled droplet environments of ∼1% size variation (Guo et al., 2012). However, previous studies have not fully leveraged the capabilities of this technology to quantitatively investigate microbial communities. Further, we lack a systematic method to rapidly infer microbial interactions using droplet microfluidics in different environmental contexts.

To address this challenge, we developed <u>M</u>icrobial <u>I</u>nteraction <u>N</u>etwork Inference in <u>Drop</u>lets (MINI-Drop). To determine the absolute abundance of each strain across hundreds to thousands of samples, we developed an automated computational method coupled to fluorescence microscopy to rapidly segment droplet images and accurately count fluorescently labeled cells within each droplet. We tested the capability of MINI-Drop to accurately infer microbial interactions using a microbial interaction toolbox composed of positive and negative interactions mediated by distinct molecular mechanisms. Our results demonstrate that MINI-Drop can accurately decipher pairwise as well as higher-order interactions by analyzing droplets containing 1-3 strains. We investigated how the molecular composition of the environment shapes the ecological network of a three-member consortium. A probabilistic model of cell growth modified by microbial interactions and cell death described the variability in community states across droplets containing the same initial strain composition, providing insight into the forces shaping community assembly in small populations.

## RESULTS

### Inferring microbial interactions in microfluidic droplets

Microbial interactions represent the net impact (positive, negative or negligible) of an organism on the growth of another over a specified time interval (Cao et al., 2018). Microbial interactions can be quantified by evaluating the difference in phenotype (e.g. growth response or metabolic activity) of an organism in the absence and presence of another strain (partner strain). Encapsulation of cells in a microbial community into droplets using techniques from droplet-microfluidics enables parallel culturing of many sub-communities (**Fig. 1a**). To infer microbial interactions, we needed a scalable method to determine the absolute abundance of each strain within each droplet. The average fluorescence in each droplet was not proportional to the number of cells due to variability in cellular growth rates, which dictates the rate of dilution of the fluorescent reporter (**Fig. S1a**). Therefore, we developed an automated procedure using techniques from computer vision to rapidly identify droplets (**Fig. S1b**) and count the number of fluorescently labeled cells in each droplet (**Fig. S1c**). The droplets were binned according to strain composition (**Fig. S1d)** and the cell counts were used to infer interaction type (positive, negative or negligible), strength and directionality (see Materials and Methods).

**Fig. 1.**
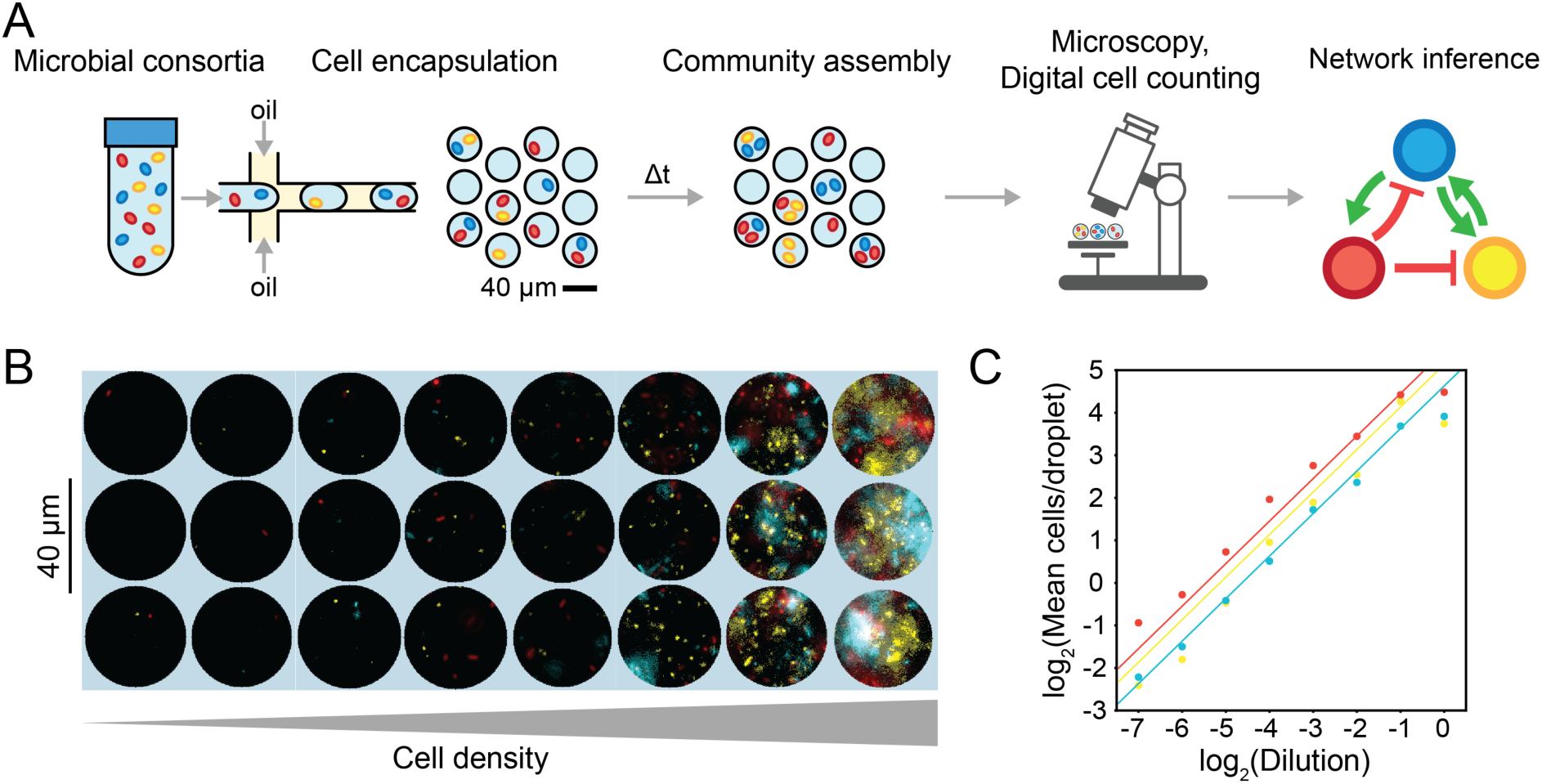
Overview and characterization of microbial interaction network inference in microdroplets (MINI-Drop). **(a)** Overview schematic of the MINI-Drop method. A mixed microbial culture and oil are loaded into a droplet-forming microfluidic device. Cells are randomly encapsulated into droplets based on a Poisson distribution. The droplets are incubated for a period of time to allow cell growth and division and then imaged using fluorescent microscopy. A computer vision workflow rapidly identifies droplets and determines the number of each fluorescently labeled strain within each droplet (**Fig. S1**). A microbial interaction network is inferred based on the difference in the mean number of cells in the absence and presence of a partner strain. **(b)** Representative fluorescent microscopy images of droplets containing three bacterial strains labeled with YFP (ST Lac*), RFP (EC WT) or CFP (EC Met-) (see **Table 2**). **(c)** Scatter plot of the dilution factor of the mixed culture vs. the log_2_ transform of the mean number of cells per drop (**Fig. S2a**). Each data point represents the mean of 400-600 droplets and lines denote linear regression fits to the data. Red, yellow and blue data points correspond to EC WT, ST Lac* and EC Met-, respectively.

To evaluate the accuracy and dynamic range of the method, CFP-labeled *E. coli*, RFP-labeled *E. coli* and YFP-labeled *S. typhimurium* were mixed in equal volumetric ratios and serially diluted to generate a broad range of cell densities (**Fig. 1b**). Each dilution of the mixed culture was encapsulated into 34 picoliter (pL) droplets (40 µm diameter), imaged using fluorescence microscopy, and analyzed using a computational workflow (see Materials and Methods). The number of cells of each fluorescently labeled strain decreased linearly with each dilution, with the exception of the highest density droplets (**Fig. 1c, Fig. S2a**). These data demonstrate at least a 64-fold linear range of the cell counting method of each fluorescent reporter. In a separate experiment involving growth of fluorescently labeled strains in droplets described below (**Table 1**, E6), droplet size did not correlate with the number of cells labeled with CFP, YFP or RFP, indicating that variation in droplet size did not contribute to variability in cell growth (**Fig. S2b,c,d**).

**Table 1.** Strains used in growth experiments.

| Number | Strains | Media | Figure |
| --- | --- | --- | --- |
| E1 | BS Trp-EC Met- (RFP) | M9 Glucose | 2a-d |
| E2 | EC Sender<br>EC Receiver | LB | 2e-h |
| E3 | EC Met- (CFP)<br>ST Lac*<br>EC WT | M9 Lactose | 3a,e,i<br>4d,e |
| E4 | EC Met- (CFP)<br>ST Lac*<br>EC WT | M9 Lactose + Met | 3b,f,j |
| E5 | EC Met- (CFP)<br>ST Lac*<br>EC WT | M9 Glucose | 3c,g,k |
| E6 | EC Met- (CFP)<br>ST Lac*<br>EC WT | M9 Glucose + Met | 3d,h,l |
| E7 | EC Met- (CFP)<br>ST Lac*<br>EC Met- Lac* (RFP) | M9 Lactose | 4b,c |

**Table 2.** Strains and media conditions for each experiment.

| Strain | Genotype | Plasmid | Fluorescent reporter | Abbreviation | Reference |
| --- | --- | --- | --- | --- | --- |
| *B. subtilis* | *B. subtilis* 168,<br>*trpC2*, *cat*,<br>*amyE::Pveg-gfp-spec* | None | GFP | BS Trp- | (Burkholder and Giles, 1947) |
| *E. coli* | *E. coli* BW25113<br>*pheA::Kan* | pOSV005 | RFP | EC Met- Lac*<br>and EC Met-<br>(RFP) | (Baba et al., 2006) |
| *E. coli* | *E. coli* BW27783 | pOSV022 | GFP | Sender | (Khlebnikov et al., 2001) |
| *E. coli* | *E. coli* MG1655z1 | pOSV151 | RFP | Receiver | (Cox et al., 2007) |
| *E. coli* | *E. coli* K12<br>BW25113,<br>$\Delta metB$ att::pLC2<br>80 [kan P<sub>L'</sub>-cfp<br>oriR6K] | None | CFP | EC Met- (CFP) | (Adamowicz et al., 2018) |
| *S. typhimurium* | LT2,<br>*metA*(P35L) *met* | None | YFP | ST Lac* | (Adamowicz et al., 2018; |
|  | *J*(16:IS10) *att::p*<br>LC246 [kan<br>P<sub>L'</sub>-yfp oriR6K] |  |  |  | Douglas et al.,<br>2016) |
| *E. coli* | *E. coli* BW25113<br>metA::Kan | pOSV006 | RFP | EC WT | (Baba et al.,<br>2006) |

### Investigating microbial interaction networks two-member consortia

To determine whether MINI-Drop could illuminate microbial interactions in microbial consortia, we investigated two-member consortia engineered to display defined interactions. A microbial interaction was defined as a statistically significant difference in the average number of cells of a given strain in the presence of a second strain (partner) compared to the absence of the partner at a specific time point. To investigate positive interaction networks with MINI-Drop, we constructed a consortium composed of an RFP-labeled *E. coli* methionine auxotroph (EC Met-) and a GFP-labeled *B. subtilis* tryptophan auxotroph (BS Trp-, **Table 1**, E1). In the absence of supplemented amino acids, the growth of *B. subtilis* requires secretion of tryptophan from *E. coli* and the growth of *E. coli* requires secretion of methionine from *B. subtilis*, which together generates a bidirectional positive interaction network (**Fig. 2a**). The two species were mixed in equal proportions based on OD600 measurements, encapsulated into droplets such that each droplet had 1-2 cells on average according to a Poisson distribution and the droplets were incubated at 37°C for 18 hours. The fluorescence microscopy images demonstrated that single species droplets exhibited a low number of total cells, whereas droplets containing both species exhibited significantly higher number of cells of each strain (**Fig. 2b**). Specifically, the average number of EC Met-cells was 3.3-fold (p = 3.8e-26) higher in the presence of BS Trp-compared to the average number of EC Met-in single-species droplets (**Fig. 2c**). Similarly, the average number of BS Trp-cells was 4.2-fold (p = 1.5e-6) higher in the presence of EC Met-compared to the average number of BS Trp-cells in single-species droplets (**Fig. 2c**). The inferred interaction network exhibited bidirectional positive interactions, mirroring the topology of the expected interaction network (**Fig. 2a,d**), demonstrating that MINI-Drop could deduce positive interactions. Corroborating this result, the cell counts for BS Trp-and EC Met-were positively correlated (**Fig. S3a**).

**Fig. 2.**
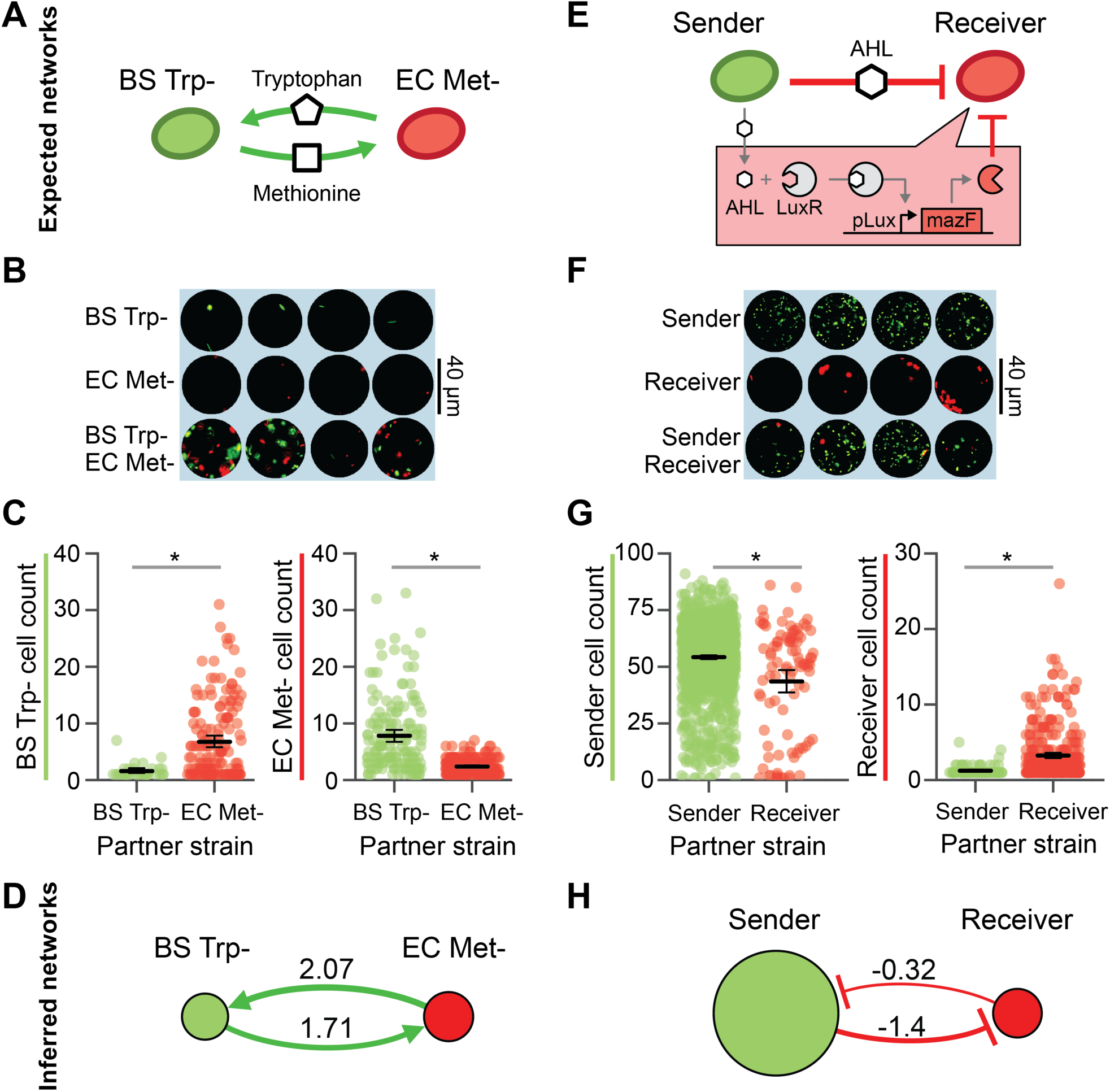
Investigating positive and negative microbial interaction networks using MINI-Drop. **(a)** Schematic of the expected network for a synthetic consortium composed of an RFP-labeled *E. coli* methionine auxotroph (EC Met-) and a GFP-labeled *B. subtilis* tryptophan auxotroph (BS Trp-) (**Table 1**, E1). **(b)** Fluorescence microscopy image of representative single-species (EC Met-or BS Trp-) or two-member droplets. **(c)** Categorical scatter plot showing the number of BS Trp-or EC Met-cells in each droplet. The black horizontal line represents the mean and the error bars denote bootstrapped 95% confidence intervals for the mean. Gray lines denote statistically significant difference in means based on the Mann-Whitney U test (n=87, p=1.5e-6, left and n=372, p=3.8e-26, right). **(d)** The inferred interaction network for the EC Met-, BS Trp-consortium. The edge width is proportional to the log_2_ ratio of the average cell count in the presence of a partner to the average cell count in single strain droplets. Node size is proportional to the average cell count of each strain in single strain droplets. **(e)** Schematic of the expected network of an *E. coli* community that exhibits a strong unidirectional negative interaction. A GFP-labeled strain (sender) expresses LuxI, a synthetase for the quorum-sensing signal C6 acyl homoserine lactone (AHL). AHL binds to the receptor LuxR in an RFP-labeled strain (receiver) and activates the expression of a toxin MazF, generating a strong negative interaction (**Table 1**, E2). **(f)** Fluorescence microscopy image of representative droplets containing the sender strain, receiver strain or community. **(g)** Categorical scatter plot of the number of sender or receiver cells in each droplet in the presence or absence of a partner. The black line represents the mean and the error bars denote bootstrapped 95% confidence intervals for the mean. Gray lines denote statistically significant differences in the means (n=1512, p=2.2e-4, left, n=421, p=3.8e-14, right). **(h)** The inferred interaction network for the mazF inhibition consortium.

We next investigated whether MINI-Drop could decipher negative interactions. A synthetic community was constructed wherein a GFP-labeled *E. coli* strain (sender strain) was engineered to express LuxI, a synthetase for the quorum-sensing signal C6 acyl homoserine lactone (AHL). AHL diffuses into the RFP-labeled *E. coli* strain (receiver strain), binds and activates the receptor LuxR, which regulates the expression of the MazF toxin (**Fig. 2e**, **Table 1**, E2). High expression levels of the endoribonuclease MazF inhibits cell growth by inducing mRNA decay (Venturelli et al., 2017), generating a strong negative interaction from the sender to the receiver. To characterize this community using MINI-Drop, the sender and receiver strains were mixed in equal proportions based on OD600, encapsulated into droplets and incubated at 37°C for 18 hr. Computational analysis of the fluorescent microscopy images showed that the number of receiver cells was significantly lower in droplets containing both the sender and receiver strains compared to the average number of receiver cells in single-strain droplets (**Fig. 2f**). The average number of receiver cells in the presence of the sender was 2.6-fold lower (p = 3.7e-14) compared to the average number of receiver cells in droplets containing the receiver strain alone (**Fig. 2g**). In addition, the average number of sender cells was 1.25-fold lower (p = 2.2e-4) in the presence of the receiver compared to in its absence (**Fig. 2g**). The average number of sender cells in droplets containing the sender strain alone was 16.7-fold higher than the average number of receiver cells in droplets containing only the receiver strain, presumably due to leakiness of *mazF* from the pLux promoter in the absence of AHL. Based on these data, the inferred interaction network exhibited a strong negative interaction from the sender to the receiver and a weak negative interaction from the receiver to the sender (**Fig. 2h**). The cell counts of the sender and receiver were negatively correlated across droplets, corroborating the presence of negative interactions (**Fig. S3b**). In sum, these data demonstrate that MINI-Drop can decipher negative interactions in microbial consortia.

### The molecular composition of the environment shapes a microbial interaction network

The molecular composition of the environment influences the energetic costs and benefits of microbial interactions in microbial communities (Cao et al., 2018; Harcombe et al., 2016; Liu et al., 2017). A key challenge is predicting how the microbial interaction network is modulated by environmental parameters. To investigate this question, we constructed a three-member community consisting of two strains interacting via bidirectional positive interactions and a third strain that promoted growth of constituent members of the community but did not receive a benefit from the community. Specifically, the strains included RFP-labeled *E. coli* (EC WT), CFP-labeled *E. coli* methionine auxotroph (EC Met-), and YFP-labeled *S. typhimurium* (ST Lac-). This consortium was characterized in four conditions that varied the carbon source (lactose or glucose) and the presence or absence of supplemented methionine. In lactose minimal media, *E. coli* can consume lactose and secrete carbon byproducts that can be utilized as substrates by ST Lac* (**Table 1**, E3-6) (Harcombe, 2010). In the absence of supplemented methionine, the growth of EC Met-is dependent on methionine provided by constituent community members.

We used MINI-Drop to infer the pairwise microbial interaction network based on the patterns in the number of cells of each community member in single strain and two-member droplets. In lactose minimal media lacking supplemented methionine, the inferred network recapitulated the expected network, exhibiting bidirectional positive interactions between ST Lac* and EC Met-and unidirectional positive interactions from EC WT to ST Lac* to EC Met-(**Fig. 3a,e,i, Table 1**, E3, **Table S1**). In lactose minimal media supplemented with methionine, the positive outgoing interactions from EC WT or ST Lac* to EC Met-were absent in the network and bidirectional negative interactions linked EC Met-and EC WT (**Fig. 3b,f,j, Table 1**, E4). In glucose minimal media lacking supplemented methionine, the positive interactions from EC WT or EC Met-to ST Lac* were absent and instead EC WT and ST Lac* were coupled by bidirectional negative interactions (**Fig. 3c,g,k, Table 1**, E5). By contrast to the expected network, bidirectional negative interactions were inferred between all pairs of strains in glucose minimal media supplemented with methionine (**Fig. 3d,h,l, Table 1**, E6). Across all conditions, the sign of the Pearson correlation coefficient clustered according to the pairwise network topology, wherein positive or negative correlation coefficients were associated with positive or negative interactions, respectively (**Fig. S3, Fig. S4**). These data show that correlations in the absolute abundance of strains across droplets can be used to classify specific topologies of two-member microbial interaction networks. Media containing lactose as a primary carbon source promoted strain co-existence in three-member droplets, suggesting that positive interactions from EC Met-or EC WT to ST Lac* are critical interactions that promote community stability across different environments (**Fig. S5a**). In sum, our results demonstrate that the microbial interaction network is highly context-dependent and the network topology changes as a function of the molecular composition of the environment.

**Fig. 3.**
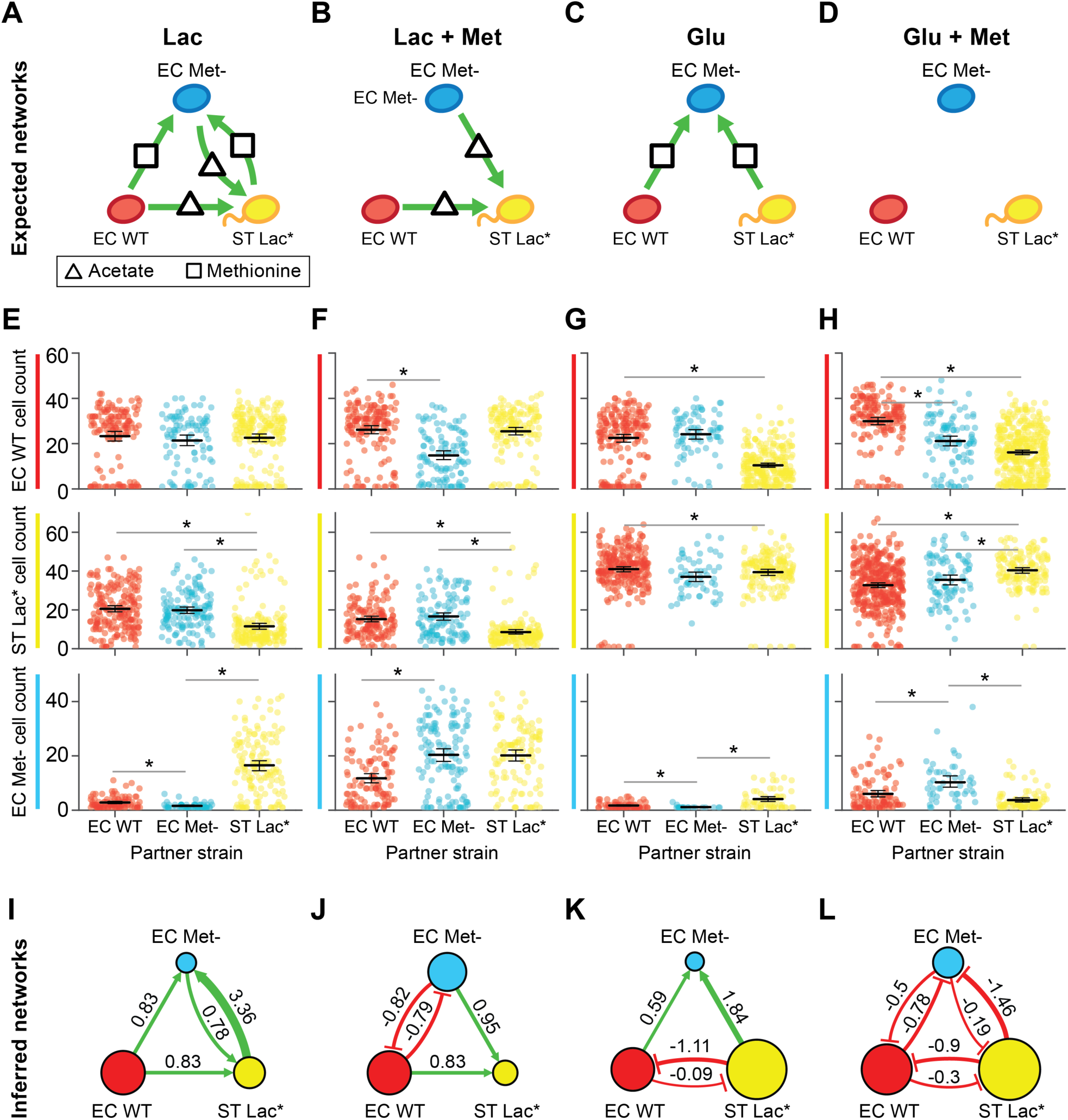
The molecular composition of the environment shapes the interaction network of a three-member consortium. **(a)** Schematic of the expected microbial interaction network of a three-member consortium consisting of RFP-labeled *E. coli* (EC WT), CFP-labeled *E. coli* methionine auxotroph (EC Met-), and YFP-labeled *S. Typhimurium* deficient in lactose metabolism (ST Lac*) in lactose minimal media lacking supplemented methionine (**Table 1,** E3). Secreted carbon byproducts (acetate) and methionine are represented by a triangle and rectangle, respectively. Node colors and green arrows denote the type of fluorescent reporter and positive interactions, respectively. **(b)** Schematic of the expected microbial interaction network in lactose minimal media supplemented with methionine (**Table 1**, E4). **(c)** Schematic of the expected microbial interaction network in glucose minimal media lacking supplemented methionine (**Table 1**, E5). **(d)** Schematic of the expected microbial interaction network in glucose minimal media supplemented with methionine (**Table 1**, E6). **(e)** Cell count distributions in lactose minimal media for EC WT (top), ST Lac* (middle) or EC Met-(bottom). The black line represents the mean and the error bars denote the bootstrapped 95% confidence intervals for the mean. The gray horizontal bars indicate a statistically significant difference (p < 0.05, **Table S1**) based on the Mann-Whitney U test. **(f)** Cell count distributions in lactose minimal media supplemented with methionine for EC WT (top), ST Lac* (middle) or EC Met-(bottom). **(g)** Cell count distributions in glucose minimal media for EC WT (top), ST Lac* (middle) or EC Met-(bottom). **(h)** Cell count distributions of EC WT (top), ST Lac* (middle) or EC Met-(bottom) in glucose minimal media supplemented with methionine. **(i)** Inferred interaction network in lactose minimal media lacking supplemented methionine. The edge width is proportional to the log_2_ ratio of the average cell count in the presence of a partner to the average cell count in the absence of the partner. Node size is proportional to the average cell count of each strain grown in isolation. **(j)** Inferred network in lactose minimal media supplemented with methionine. **(k)** Inferred interaction network in glucose minimal media lacking supplemented methionine. **(l)** Inferred interaction network in glucose minimal media supplemented with methionine.

### Investigating higher-order interactions using MINI-Drop

Higher-order interactions occur when a pairwise interaction is modified in the presence of a third community member (Bairey et al., 2016; Society, 2015) and these interactions are challenging to identify in microbial communities. In MINI-Drop, a higher-order interaction was defined as a difference in the presence and sign (positive or negative) of an interaction in a three-member community compared to the presence and sign of the interaction in each two-member sub-community (**Fig. 4a**). We tested whether MINI-Drop could identify higher-order interactions by analyzing the cell count distributions of each strain in three-member droplets in addition to single-strain and two-member droplets. To do so, a community consisting of an RFP-labeled *E. coli* methionine auxotroph that is also deficient in lactose metabolism (EC Met-Lac*, **Table 1**, E7), EC Met-(CFP) and ST Lac* was constructed. In lactose minimal media lacking supplemented methionine, EC Met-and ST Lac* can secrete carbon byproducts and methionine, respectively and thus together enable the growth of EC Met-Lac*. Our results showed that the number of EC Met-Lac* cells was higher in the presence of both EC Met-and ST Lac* but not in the presence of either single strain, demonstrating that MINI-Drop could identify higher-order interactions (**Fig. 4b**, p=0.0012). The strains EC Met-(CFP) and ST Lac* interacted via bidirectional positive interactions, recapitulating the expected network topology (**Fig. 3a**, **Fig. S5b,c**). In addition, the cell counts of EC Met-and ST Lac* displayed a strong positive correlation (**Fig. S3d**).

**Fig. 4.**
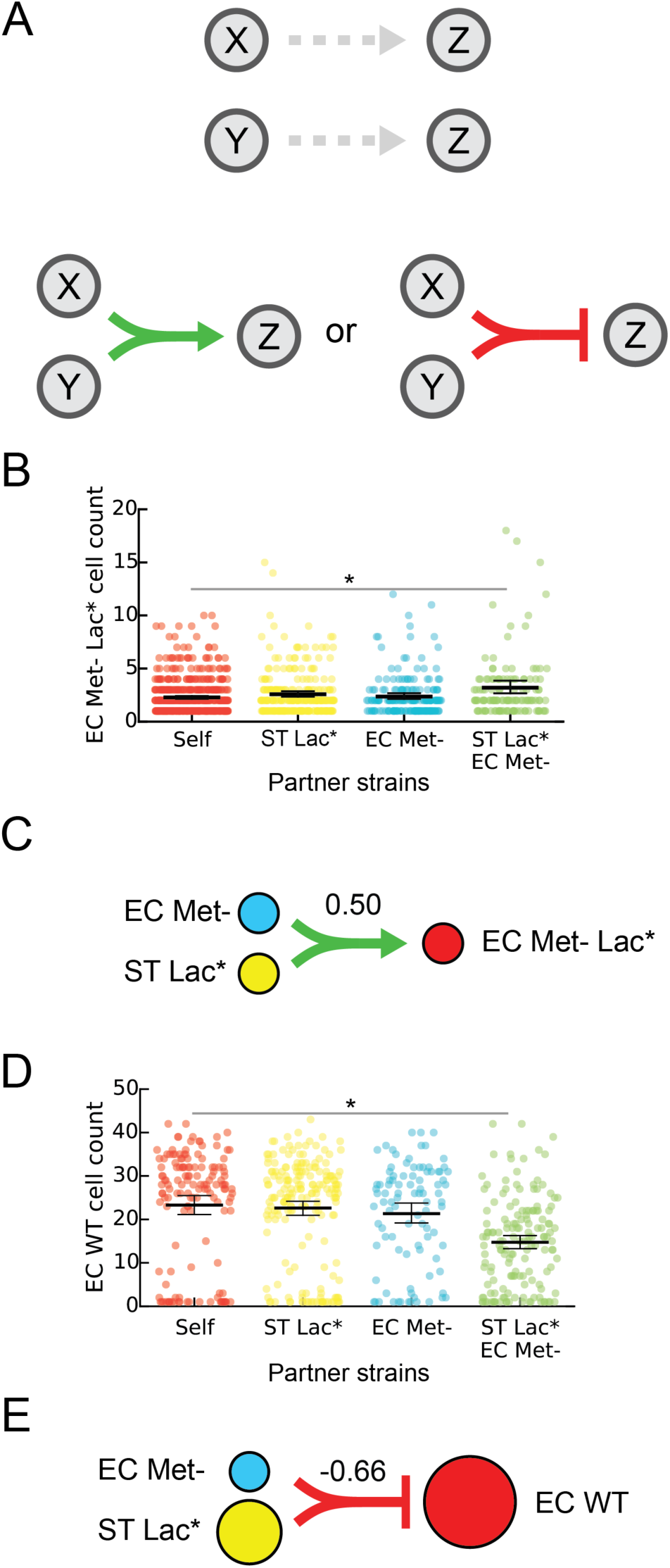
Investigating higher-order interactions using MINI-Drop. **(a)** Schematic showing an example of a higher-order interaction. Droplets containing two strains X and Z or Y and Z do not exhibit interactions. In three-member droplets, a negative or positive interaction from X and Y to Z is present and is defined as a higher-order interaction. **(b)** Categorical scatter plots of the number of EC Met-Lac* cells in droplets containing the single strain EC Met-Lac* (self), pairs of strains including EC Met-Lac* and EC Met-or ST Lac* or all three strains (EC Met-Lac*, EC Met-and ST Lac*). Black horizontal bars denote the mean number of cells per droplet and error bars represent the bootstrapped 95% confidence interval for the mean. The horizontal bar (gray) represents a statistically significant difference in means based on the Mann-Whitney U test (p = 1.2e-3, n = 703). **(c)** Schematic showing the higher-order inferred network for the data shown in panel (b). The line width represents the inferred strength of the higher-order interaction. Node size is proportional to the average cell count of each strain grown in isolation. **(d)** Categorical scatter plots of the number of EC WT cells in droplets containing the single strain EC WT, two strains including EC WT and ST Lac* or EC Met-or all three strains (EC WT, ST Lac* and EC Met-) in lactose minimal media. The horizontal bar (gray) represents a statistically significant difference in means based on the Mann-Whitney U test (p = 2.9e-10, n = 296). **(e)** Schematic showing a higher-order interaction inferred using the data shown in (d). The line width represents the strength of the inferred higher-order interaction. Node size is proportional to the average cell count of each strain grown in isolation.

To investigate other higher-order interactions that were present in our data, we analyzed droplets containing three-member consortium (EC WT, EC Met-and ST Lac), two-member sub-communities and single strains across four different environments (**Fig. 3, Table 1**, E3-E7). Our results illuminated a higher-order interaction in lactose minimal media (**Table 1**, E3), where EC WT was significantly inhibited in the presence of both EC Met-and ST Lac*, while no negative interaction was observed in the pairwise interaction networks of EC WT co-cultured with EC Met-or ST Lac* (**Fig. 3a, 4d,e**). Unexpected higher-order interactions occurred in one of twelve possible cases (3 community members times 4 environments) in the EC Met-, EC WT, ST Lac* consortium (**Table 1**, E3-6), In sum, our results show that MINI-Drop can rapidly elucidate higher-order interactions based on the absolute abundance patterns in droplets containing 1-3 strains. demonstrating that higher-order interactions were infrequent in this community across different environmental conditions.

To evaluate the sensitivity of the method, we next investigated the number of droplets containing the same initial strain composition (replicates) required to infer microbial interactions of different strengths across all datasets. Specifically, we analyzed the relationship between interaction strength magnitude, number of replicates, and interaction significance (p<0.05) in all datasets (**Fig. S6**). Our results showed that the significance of each interaction increased exponentially as a function of the number of droplets (**Fig. S6a**). The strength of the interaction was inversely related to the number of droplets required for statistical significance of the interaction. For example, strong interactions required as few as 15 replicates whereas weak interactions required more than 50 replicates (**Fig. S6b**).

### Discrete-time Markov model of community assembly

In small microbial populations, stochastic variation in intracellular molecular concentrations, growth and death can impact community assembly and functions (Boedicker et al., 2009; Connell et al., 2014; Hansen et al., 2016). To model community assembly in small populations, microbial growth can be represented as a probabilistic event, such that two communities seeded with the same initial strain composition exhibit different steady-state community compositions (**Fig. 5a**) (Horowitz et al., 2010). We constructed a discrete-time Markov model of cell growth modified by microbial interactions to investigate the variability in community composition across droplets containing the same initial strain composition.

**Fig. 5.**
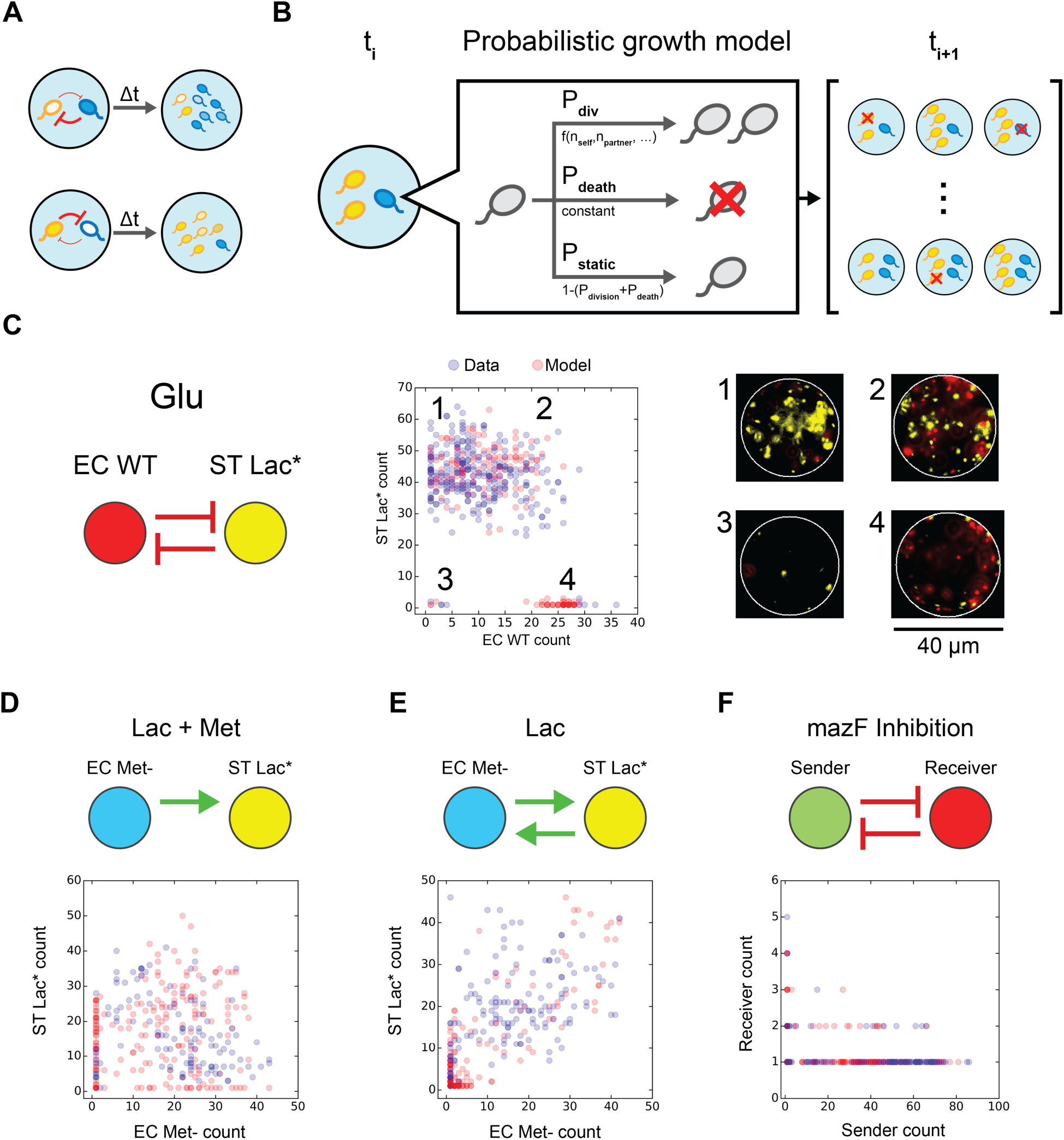
Discrete-time Markov model of cell growth modified by microbial interactions can recapitulate cell count distributions in microfluidic droplets. **(a)** Schematic of variability in community assembly in small populations. Stochasticity in intracellular molecular concentrations can alter the strength of microbial interactions, generating different community states (high blue cells, low yellow cells or the reciprocal). **(b)** Schematic of the discrete-time Markov model of cell growth modified by microbial interactions. At each time step, each cell can undergo cell division, cell death or remain static according to the probabilities P_div_, P_death_ or P_static_, respectively. **(c)** Inferred network topology using MINI-Drop (left) for the EC WT, ST Lac* consortium in glucose minimal media (**Table 1**, E5). Node size and edge weight represent the average cell count of each strain grown in isolation and the interaction strength, respectively. Scatter plot of experimentally measured cell counts (blue circles, n=257) of EC WT and ST Lac* or model steady-states (red circles, n=200). This bidirectional negative interaction network generated qualitatively different community compositions corresponding to (1) low and high EC WT and ST Lac*, respectively, (2) high EC WT and ST Lac*, (3) low EC WT and ST Lac*, (4) high EC WT and low ST Lac*. Fluorescence microscopy images (right) of a representative droplet in each community state 1-4 are shown (right). **(d)** Inferred network for the EC Met-, ST Lac* consortium (top) in lactose minimal media supplemented with methionine (**Table 1**, E4). Scatter plot of experimentally measured cell counts (blue circles, n=118) of EC Met-and ST Lac* or model steady-states (red circles, n=200). **(e)** Inferred interaction network for the EC Met-, ST Lac* consortium in lactose minimal media (top, **Table 1**, E3). Scatter plot of experimentally measured cell counts (blue circles, n=141) of EC Met-and ST Lac* or model steady-states (red circles, n=200). **(f)** Inferred interaction network for the sender, receiver consortium (top, **Table 1**, E2). Scatter plot of experimentally measured cell counts (blue circles, n=93) of the sender and receiver strains or model steady-states (red circles, n=200).

In the model, communities are initially seeded with a single cell of each type. At each time step, strain *i* can undergo cell division, death or remain static according to the probabilities *P*_*div,i*_, *P*_*death,i*_, and *P*_*static,i*_, respectively (**Fig. 5b**). The probabilities *P*_*div,i*_, and *P*_*static,i*_, are a function of the number of cells of each strain with parameters specific to each strain and the probability *P*_*death,i*_, is a fixed parameter. Negative interactions with self or non-self are represented by inverted sigmoidal logistic functions, such that the probability of cell division is inversely related to the cell number. Positive interactions are represented as sigmoidal logistic functions, such that the probability of cell division increases as a function of the number of partner cells (see Materials and Methods).

We tested whether this modeling framework could recapitulate the experimental cell count distributions, based on the assumption that the measurement time point maps to the steady-state of the model. Models were constructed using the positive or negative interaction functions and model parameters were identified to recapitulate the cell count distributions of each strain. We constructed a model for the EC WT, ST Lac* community grown in glucose minimal media that exhibited a bidirectional negative interaction network (**Fig. 5c**, left). Our results showed three clusters representing distinct community states exhibiting high abundance of one strain (**Fig. 5c**, center, clusters 1 and 2), co-existence of both strains (**Fig. 5c**, center, cluster 4), or low cell counts of both strains (**Fig. 5c**, center, cluster 3). Representative images of droplets from each cluster showed significant differences in community composition (**Fig. 5c**, right). A model of a bidirectional negative interaction network displaying strong and weak negative interactions was able to recapitulate the cell count distribution (**Fig. 5c**, middle, **Table S3**).

We next evaluated whether the model could recapitulate the cell count distributions of networks that displayed positive interactions. Models constructed for the EC Met-, ST Lac* consortium in two different environments exhibiting unidirectional or bidirectional positive interactions (**Table 1**, E3-4) could recapitulate the cell count distributions (**Fig. 5d,e**). Next, a model was developed for the mazF inhibition consortium (**Table 1**, E2) that displayed a bidirectional negative interaction network. A model of strong and weak bidirectional negative interactions represented the negative correlation in cell counts of the sender and receiver strains (**Fig. 5f**). Our results demonstrate that bidirectional negative interaction networks can realize distinct community state distributions (**Fig. 5c,f**). In the model, the number of partner cells required to impact the probability of cell division dictates the strength of an interaction (**Fig. 5f**, **Fig. S7**). The toxin mediated negative interaction in the mazF inhibition consortium (**Table 1**, E2) exhibited a higher sensitivity to partner cell number than the negative interaction from ST Lac* to EC WT in glucose minimal media (**Table 1**, E5, **Fig. S7**). Therefore, the recipients of the strong negative interactions displayed different sensitivities to variations in donor cell number, providing insight into the qualitative dissimilarities in the cell count distributions. In sum, the model was able to describe the cell count distributions for positive and negative interactions mediated by distinct molecular mechanisms, illustrating that a probabilistic growth model can explain the variability in community states in small populations.

## Discussion

We showed that MINI-Drop can rapidly infer pairwise as well as higher-order microbial interactions in 2-to 3-member consortia in different environmental conditions. This method can be scaled to quantify interactions in higher-dimensional (>3 members) communities using compatible fluorescent labels or combinatorial fluorescent imaging of multiple reporters within the same cell. Combinatorial labeling via fluorescence in situ hybridization (Liu et al., 2011; Valm et al., 2012) or fluorescent labeling of the bacterial outer membranes via biorthogonal click chemistry could be used to measure the absolute abundance of organisms that are not genetically tractable (Geva-Zatorsky et al., 2015).

In MINI-Drop, a single experiment generates hundreds to thousands of replicates of unique sub-communities. The initial mean number of cells per drop can be manipulated to investigate the contribution of initial cell density to microbial interactions or increase the proportion of multi-strain droplets for interrogation of higher-order interactions. MINI-Drop does not require coexistence of community members to determine interactions because fluorescently tagged single cells encapsulated in a droplet can be accurately quantified.

Previous methods of microbial interaction inference using modeling frameworks such as the generalized Lotka-Volterra (gLV) model are constrained by mathematical relationships (Momeni et al., 2017). For example, a gLV model of strong bidirectional positive interactions tends to be unstable, leading to potential underrepresentation of bidirectional positive interactions. Further, it is challenging to pinpoint if the failure of a pairwise gLV model to accurately fit experimental data is attributed to the presence of higher-order interactions or to unmodeled dynamics such as metabolites mediating the interactions. By contrast, MINI-Drop is not constrained to a defined mathematical framework and thus can rapidly identify higher-order interactions in the networks. We showed that MINI-Drop accurately inferred diverse interaction topologies including unidirectional positive, bidirectional positive or bidirectional negative networks. In addition to deciphering engineered interactions, MINI-Drop illuminated unexpected negative pairwise interactions and higher-order interactions in the networks. The unexpected higher-order interaction that inhibited EC WT in the presence of both EC Met-and ST Lac* could be explained by robust growth of the mutualistic pair EC Met-and ST Lac*, which in turn negatively impacted EC WT. Across all experiments, *S. typhimurium* exhibited the strongest outgoing negative interactions as well as the highest carrying capacity, suggesting that the negative interactions could arise from competition for limited resources or space or the production of toxic compounds.

The throughput of the MINI-Drop method was enabled by coupling two automated and scalable technologies, droplet microfluidics and computational image analysis. The large number of sub-community replicates provided by MINI-Drop allows investigation of the contribution of initial conditions to community assembly in small populations. A probabilistic analysis of the distribution of community states provides insight into the stochastic nature of microbial interactions and impact of these parameters on community assembly. For example, we observed that bidirectional positive networks displayed frequent co-occurrence (**Fig. 5e, Fig. S3, Fig. S4**) whereas a bidirectional negative network can realize a set of distinct community states (**Fig. 5c,e**).

Our stochastic growth model can recapitulate the community states observed experimentally a set of synthetic communities. This demonstrates that a simple probabilistic representation of cell growth, death and microbial interactions can give rise to multiple community steady-states from the same initial conditions. Our modeling framework could be used to predict the probability of strain growth as a function of the initial strain proportions and cell density. These parameters could be manipulated to maximize the likelihood of community member coexistence in multi-species consortia. In sum, we developed a systematic procedure to elucidate microbial interaction networks in microdroplets. Future work will apply MINI-Drop to study diverse cellular interactions such as interkingdom or mammalian cell interactions.

## MATERIALS AND METHODS

### Dynamic range of cell counting

The bacterial strains EC Met-(CFP), EC WT (RFP), and ST Lac* (YFP) were grown in LB medium to early stationary phase, centrifuged at 18,000xg for 1 min, decanted, and resuspended in M9 minimal medium without glucose. Next, the cells were centrifuged at 18,000xg for 1 min, decanted and resuspended in a smaller volume of M9 minimal medium without glucose to concentrate the cells. The OD600 values of the concentrated EC Met-, EC WT and ST Lac* cultures were 14.4, 19.6, and 6.4, respectively. Equal volumes of each culture were combined to generate the mixed culture. The mixed culture was serially diluted by a factor of 2 until a dilution of 2^-7^ was reached. The diluted cultures were encapsulated separately using the droplet maker device and the resulting droplets were imaged and quantified using the computational image analysis pipeline.

### Bacterial cell culturing

Strains were grown for approximately 12 hours at 37°C in LB, diluted 1:50 into fresh LB, and then grown to an OD600 of 0.3-1. Next, the culture (3 mL) was centrifuged for 2 min at 3,500 x g and supernatant was removed. The cells were washed 4X by resuspending the pellet in 0.5 mL of minimal media and centrifuged as described above. The cell cultures containing different strains were normalized to an OD600 of 0.15 and mixed in a 1:1 ratio. In the mutualism experiment (E2), *B. subtilis* and *E. coli* were mixed in a 2:1 volumetric ratio to account for differences in the cell number to OD ratios. In experiment E1, cells were cultured in M9 supplemented with glucose (1X M9 salts, 2 mM MgSO_4_, 100 µM CaCl_2_, 0.4% glucose) and 25 μg/mL chloramphenicol (Sigma). In experiment E2, cells were cultured in LB media containing 50 ng/mL anhydrotetracycline (aTc, Cayman Chemicals), 0.1% arabinose (Sigma) and 25 μg/mL chloramphenicol. In experiments E3-E7, cells were cultured in M9 media (1X M9 salts, 2 mM MgSO_4_, 100 µM CaCl_2_) supplemented with 0.4% glucose, 0.2% lactose and/or 200 µM methionine as indicated.

### Fabrication of microfluidic devices

Photoresist masters of 25 µm layer height were fabricated by spinning a layer of photoresist SU-8 3025 (Microchem) onto a silicon wafer (University Wafer), then baked at 95°C for 10 minutes. Following baking, photoresist master was patterned by UV photolithography over a photomask (**File S1**, CADArt). The master was subjected to post-exposure bake at 95°C for 4 min and developed in fresh SU-8 developer (Microchem) for 6 min, prior to rinsing with isopropyl alcohol (Fischer Scientific) and baking at 150°C to remove the solvent. The microfluidic devices were fabricated by pouring poly(dimethylsiloxane) at a 11:1 polymer-to-crosslinker ratio (Dow Corning Sylgard 184) onto the master and curing at 65°C for 1 hr. The PDMS devices were excised with a scalpel and cored with a 0.75 mm biopsy core (World Precision Instruments) to create inlets and outlets. The device was then bonded to a microscope glass slide using O2 plasma cleaner (Harrick Plasma), and channels were treated with Aquapel (PPG Industries) to render them hydrophobic. Finally, the devices were baked at 65°C for 20 min to evaporate excess Aquapel prior to use.

### Encapsulation of cells into droplets and fluorescence microscopy

To encapsulate cells into droplets, 1 mL syringes (BD Luer Lok) were fitted with 27 gauge needles and PE/2 tubing. 500 µL of the culture was loaded into a 1 mL syringe. Hydrofluoroether oil was prepared with 2% Krytox as surfactant and loaded into a 1 mL syringe. The free end of the tubing was primed and inserted into the droplet-making device. Droplets were generated using 600 µL hr^-1^ oil and 300 µL hr^-1^ cell mixture flow rates at a 30 µm x 25 µm junction, which generated ∼40 µm diameter droplets at 4.8 kHz. Droplets were collected into a 1.5 mL microfuge tube for 15 min and incubated for 18 hr at 37°C. Droplets were imaged using chamber microscopy slides (Invitrogen C10228) and imaged with a 20X objective (Nikon, MRH10201) on a Ti-E Eclipse inverted microscope (Nikon). Fluorescence was imaged using the following filters (Chroma): (1) CFP: 436nm/20nm (ex), 480nm/40nm (em); (2) GFP: 470nm/40nm (ex), 525/50nm (em); (3) RFP: 560nm/40nm (ex), 630/70nm (em); and (4) YFP: 500nm/40nm (ex), 535nm/30nm (em).

### Fluorescence microscopy image analysis

Custom code in Python was used for automated cell counting in droplets and microbial interaction network inference. Droplets were identified from the phase contrast image using the Hough transformation algorithm (OpenCV 3). Droplets with a diameter 10% larger or smaller than 40 µm were removed from the dataset. Fluorescent cells were segmented by identifying connected regions using the SimpleBlobDetector object (OpenCV 3). Droplets were binned by the presence or absence of each fluorescently labeled strain. Interaction strength from strain *j* to strain *i*, where droplet *d* contains *d*_*k*_ cells of strain *k*, was defined according to Equation 1. Network schematics were drawn with Cytoscape 3.5 (Shannon et al., 2003).

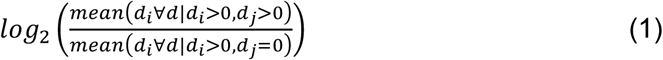

### Discrete-time Markov model of cell growth

A discrete-time Markov model was developed to recapitulate the experimentally measured cell count distributions. At each time step, the propagation of each strain is determined by computing the probability of cell division (*P*_*div,i*_,), cell death (*P*_*death,i*_,), and remaining unchanged (*P*_*static,i*_,) (Equations 2-4).

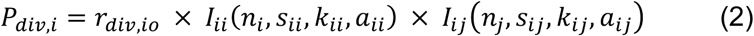

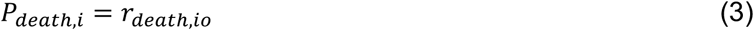

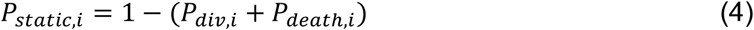

The parameter *r*_*div,io*_is the basal probability of cell division for strain *i*. The parameter *r*_*death,io*_ represents the probability of cell death of strain *i* (constant). *n*_*i*_ denotes the number of cells of strain *I* and *s*_*ij*_ defines whether the outgoing interaction of strain *j* (donor) to strain *i* is positive (*s*_*ij*_ = 1) or negative (*s*_*ij*_ = –1). The parameters *k*_*ij*_ and *a*_*ij*_ define the sigmoidal interaction function *I*_*ij,*_ representing the incoming interaction for strain *i* produced by strain *j* (Equation 5).

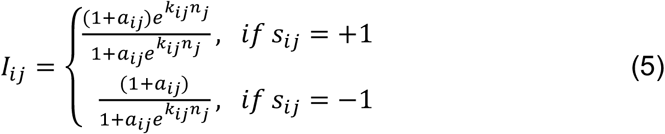

The negative interaction function approaches zero as a function of *n*_*j*_ whereas the positive interaction approaches (1 +*a*_*ij*_)/*a*_*ij*_ as a function of *n*_*j*_. The values of *a*_*ij*_ and *r*_*div,I*_ are constrained such that *P*_*div,i*_≤ 1 (Equation 6). The self-interaction function *I*_*ii*_(*n*_*i*_, *s*_*jj*_, *k*_*ii*_, *a*_*ii*_) is less than one (*s*_*ii*_ = –1) and approaches zero as a function of *n*_*i*_, leading to saturation of the number of cells of strain *i*. The interaction function *I*_*ij*_, is equal to 1 when *n*_*j*_ = 0, representing the absence of an interaction between strain *i* and *j*. In the absence of an interaction between strain *i* and *j, P*_*div,i*_ is not dependent on strain *j* (*s*_*ij*_ = –1, *k*_*ij*_ = 0, *a*_*ij*_ = 0). The outgoing interaction from the partner strain *j I*_*ij*_(*n*_*j*_, *s*_*ij*_, *k*_*ij*_, *a*_*ij*_)can be positive or negative depending on the value of the parameter *s*_*ij*_. The parameters *a*_*ij*_ and *k*_*ij*_ determine the interaction sensitivity defined as the number of partner cells at the half-maximum of the interaction function The 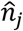 parameters

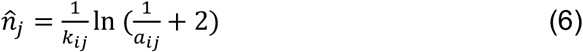

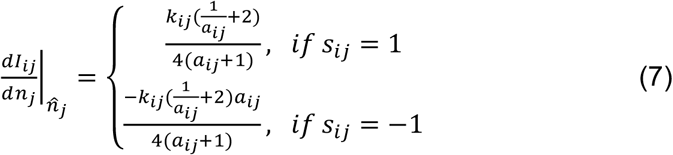

At each time step, the state transition of a cell is independent of all other cells and the cell’s prior history. The state transitions were simulated by sampling from a trinomial distribution determined by the probabilities *P*_*div,i*_, *P*_*death,i*_, and *P*_*static,i*_, Communities were simulated for 100 time-steps wherein each time-step corresponded to 10.8 minutes of experimental time. Variables were constrained such that the cell populations reached a steady state within the simulation time. The initial condition for all simulations was *n*_*i*_,*n*_*j*_ = 1. Model parameters are listed in **Table S3**.

## ACKNOWLEDGEMENTS

We would like to thank William Harcombe (University of Minnesota) for generously providing the *E. coli* Met-(CFP) and *S. typhimurium* (YFP) strains. We are grateful to Sonali Gupta and Yu-Yu Cheng for help with strain construction and Leland Hyman for assistance with droplet-microfluidics. This work was supported by the Army Research Office Young Investigator Award W911NF-17-1-0296.

## AUTHOR CONTRIBUTIONS

O.S.V., R.H.H. and P.A.R. designed the research. R.H.H., J.W.T., R.L.C. carried out the experiments. R.H.H. designed and implemented data analysis methods and computational modeling. R.H.H., R.L.C. and O.S.V. wrote the manuscript and J.W.T. assisted in revising the manuscript.

