## Supplementary Information for "Rapid microbial interaction network inference in microfluidic droplets"

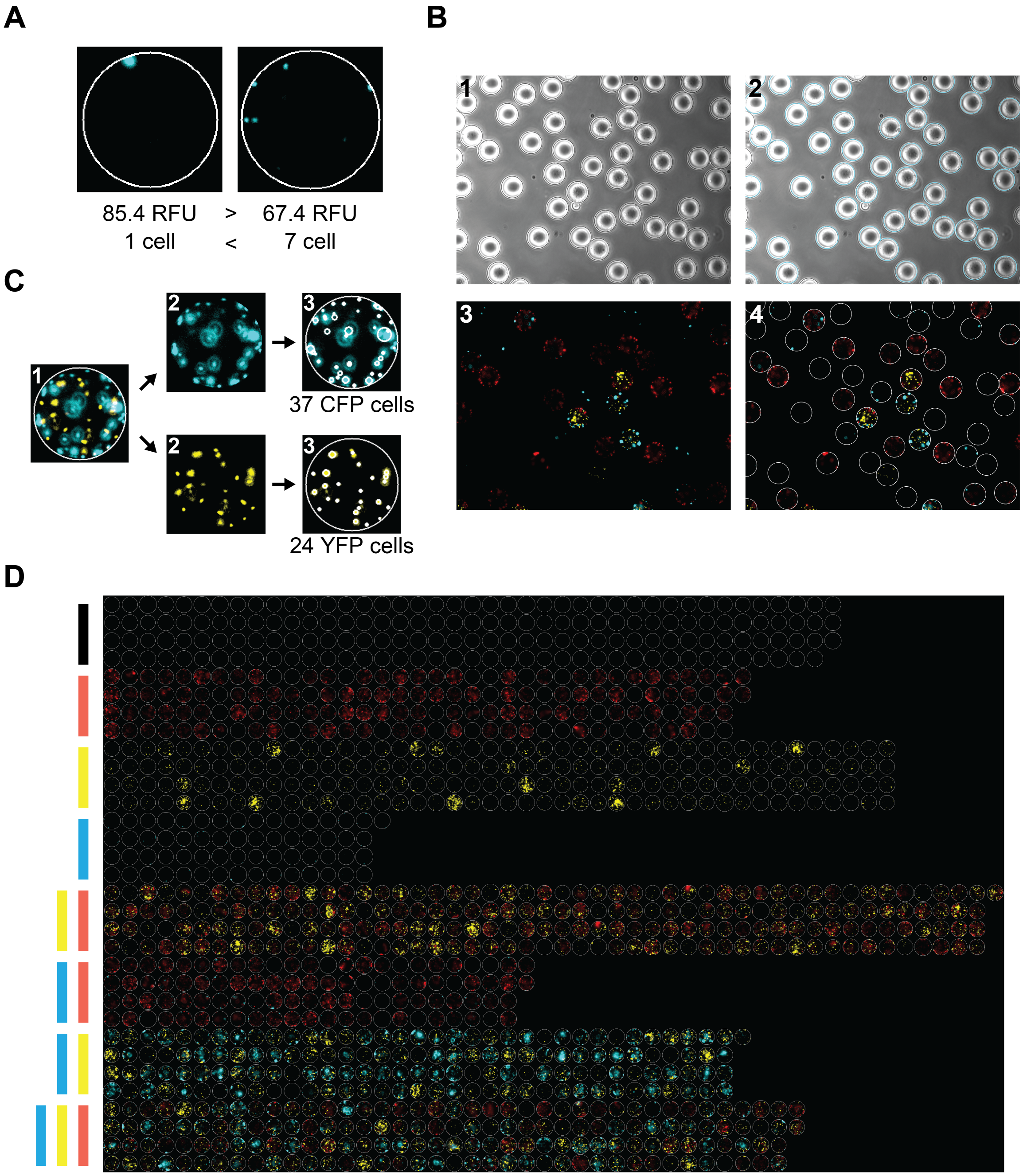


**Fig S1. Computational algorithm for droplet identification and cell counting. (a)** Quantification of strain abundance by automated cell counting or computing the average fluorescence intensity (relative fluorescence units). A representative droplet (left) contained seven times fewer cells and exhibited higher average fluorescence intensity than a different representative droplet (right). **(b)** Representative microscopy images showing the droplet segmentation pipeline. First, the raw bright field image is loaded (1) and droplet positions and size (2, blue circles) are identified via the Hough transformation (OpenCV). The identified droplet locations and raw fluorescence microscopy images (3) are used to segment the droplets (4, white circles). **(c)** Representative fluorescence microscopy images showing cell counting pipeline. Each segmented droplet image (1) is split into separate images for each fluorescent channel (2). Cells are counted by identifying connected regions of intensity using a blob detection algorithm in OpenCV (3). **(d)** Representative fluorescence microscopy image showing the final combined droplets for an experiment binned according to the presence and absence of each fluorescently labeled strain.

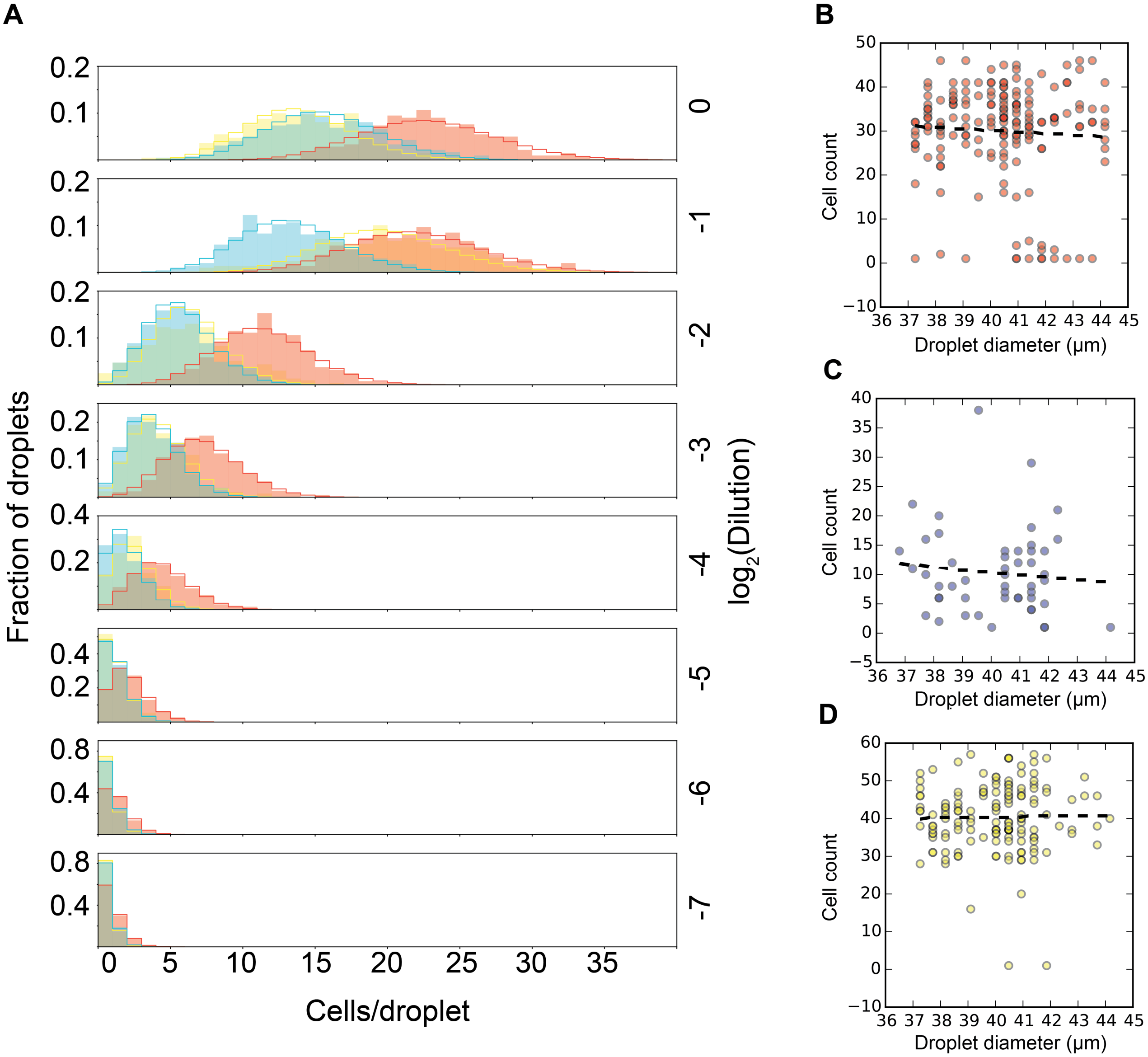


**Fig. S2. Characterization of cell counting method in microfluidic droplets. (a)** Histograms of the number of cells in each droplet for a mixed culture of CFP-labeled *E. coli* (blue), RFP-labeled *E. coli* (red) and YFP-labeled *S. typhimurium* (yellow). Each subplot represents a different dilution. Solid lines represent the expected Poisson distributions with mean (λ) equal to the mean number of counts for the corresponding distribution. **(b)** Scatter plot of droplet diameter vs. EC WT cell count (slope=-0.35, r = -0.06, n=177). Data corresponds to the three-member consortium (EC

WT, EC Met- and ST Lac*) grown in glucose minimal media supplemented with methionine (**Table 1**, E6). **(c)** Scatter plot of droplet diameter vs. EC Met- cell count (slope=-0.43, r = -0.10, n=50). **(d)** Scatter plot of droplet size vs. ST Lac* cell count (slope=0.12, r = -0.02, n=148).

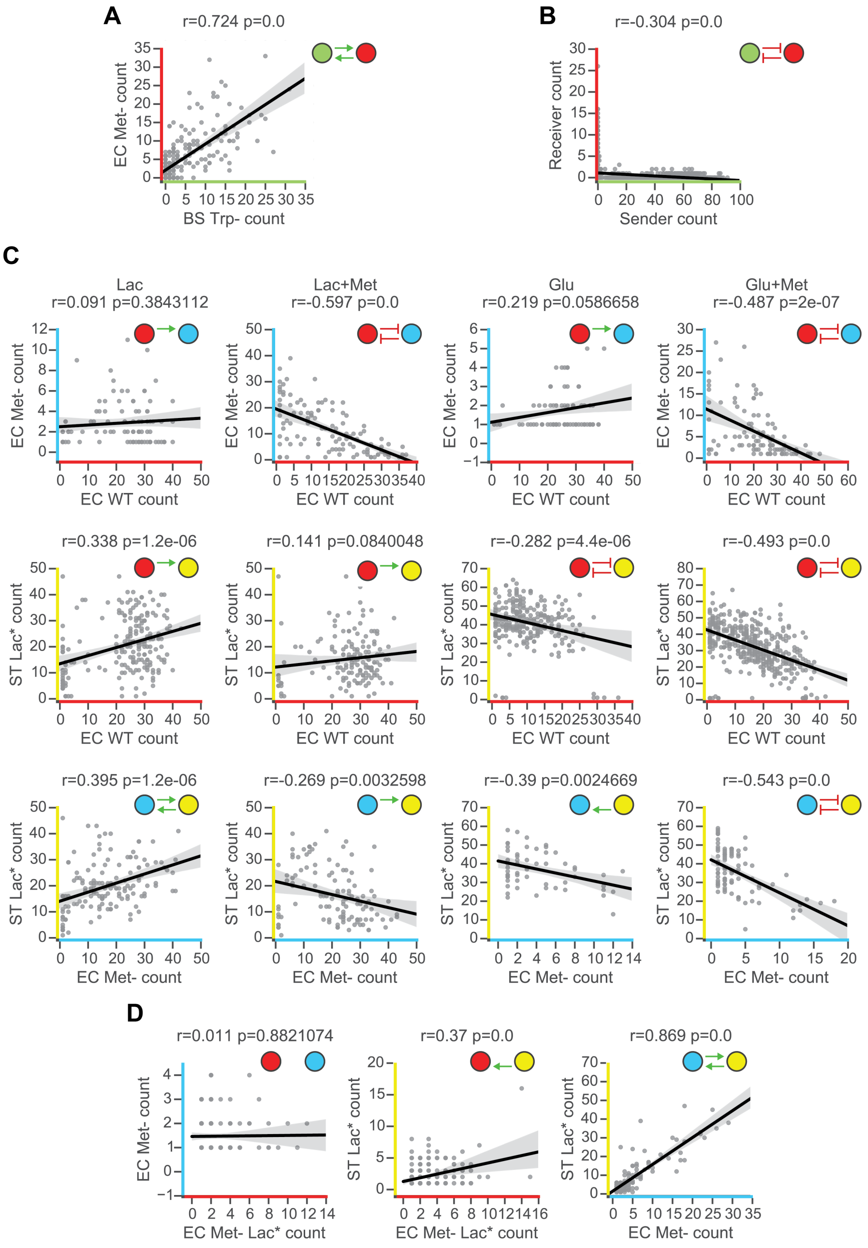


**Fig. S3. Correlation analysis of cell counts for each pair of strains across all experiments.** **(a)** Scatter plot of BS Trp- and EC Met- cell counts for experiment E1 (**Table 1**). The black line represents the regression line and the gray regions represent the bootstrapped 95% confidence interval for the regression line. Axes and nodes are colored by the fluorescent reporter of each strain and the network schematic represents the inferred interaction topology. **(b)** Scatter plot of sender and receiver *E. coli* strains in experiment E2 (**Table 1**). **(c)** Scatter plot of all pairs of strains in the three-member experiment E3-E6 (**Table I**) grown in lactose minimal media (left column), lactose minimal media supplemented with methionine (second column from the left), glucose minimal media (third column from the left) or glucose minimal media supplemented with methionine (right column). **(d)** Scatter plot of all pairs of strains in the engineered higher-order interaction experiment E7 (**Table 1**).

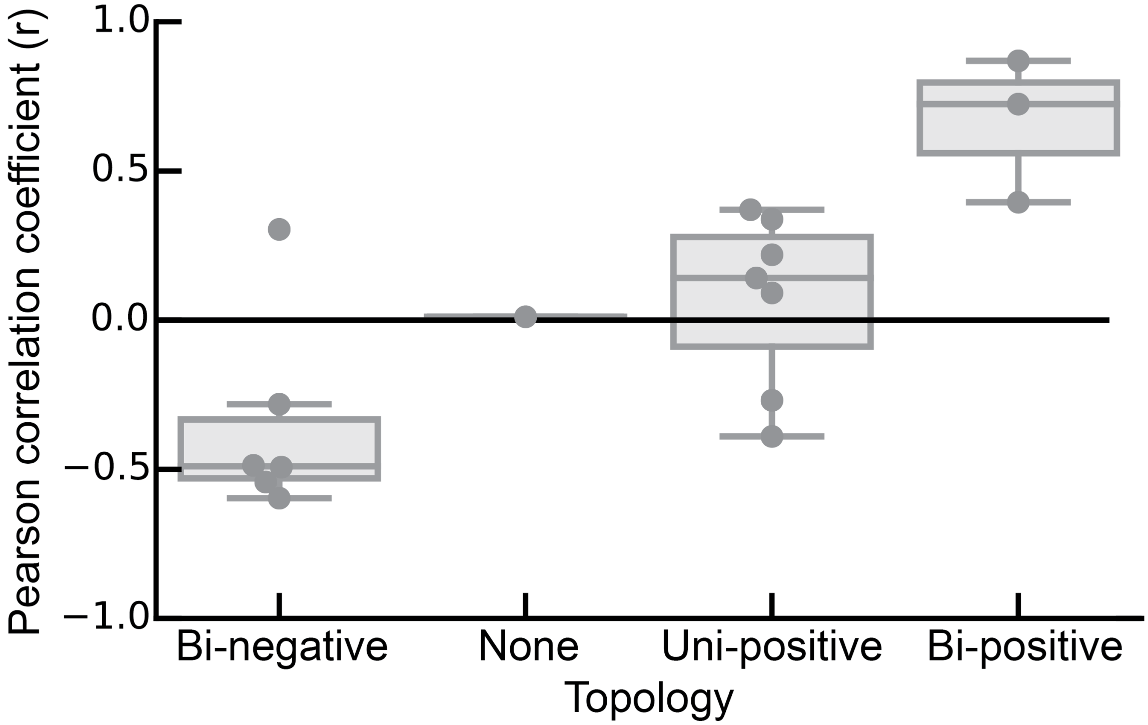


**Fig. S4. Pearson correlation coefficient clusters by pairwise network topology.** Box and whiskers plot of the Pearson Correlation coefficients of cell counts for each pair of strains across all experiments binned by the inferred network topology using MINI-Drop. The center line indicates the median, the bottom and top edges of the box indicate the 25$^{th}$ and 75$^{th}$ percentiles, respectively. The whiskers extend to the most extreme data points not considered outliers and outliers are plotted individually using the diamond symbol.

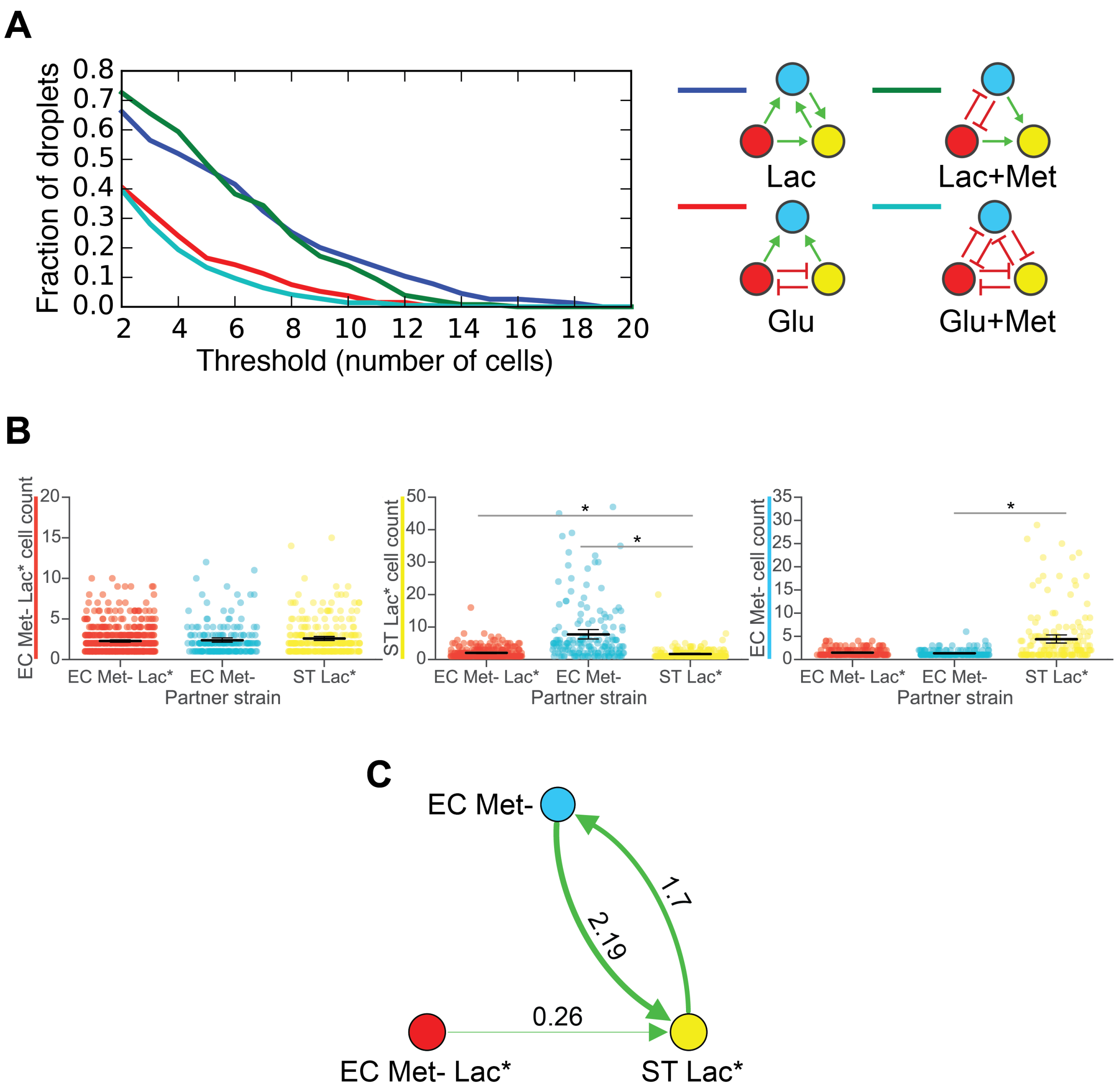


**Fig. S5. Quantification of microbial interactions and strain coexistence in three-member consortia. (a)** Fraction of three-member droplets that exhibit strain coexistence in a three-member consortium (EC Met-, EC WT, ST Lac*, **Table 1**, E3-6) following 18 hr incubation at 37°C. The plot shows the threshold number of cells (x-axis) vs. the fraction of three-member droplets. Lines of different colors represent the community grown in different media conditions. **(b)** Categorical scatter plot showing cell count distributions of EC Met- Lac* (top), ST Lac* (middle) and EC Met- (bottom) in lactose minimal media (**Table 1**, E7) The black horizontal line represents the mean and the error bars denote the bootstrapped 95% confidence intervals for the mean. The gray horizontal bars indicate a statistically significant difference ($p < 0.05$, **Table S1**). **(c)** Schematic of the inferred interaction network. The edge width is proportional to the $\log_2$ ratio of the average cell count in the presence of a partner compared to the cell count in the absence of the partner. Node size is proportional to the average cell count of each strain grown in isolation.

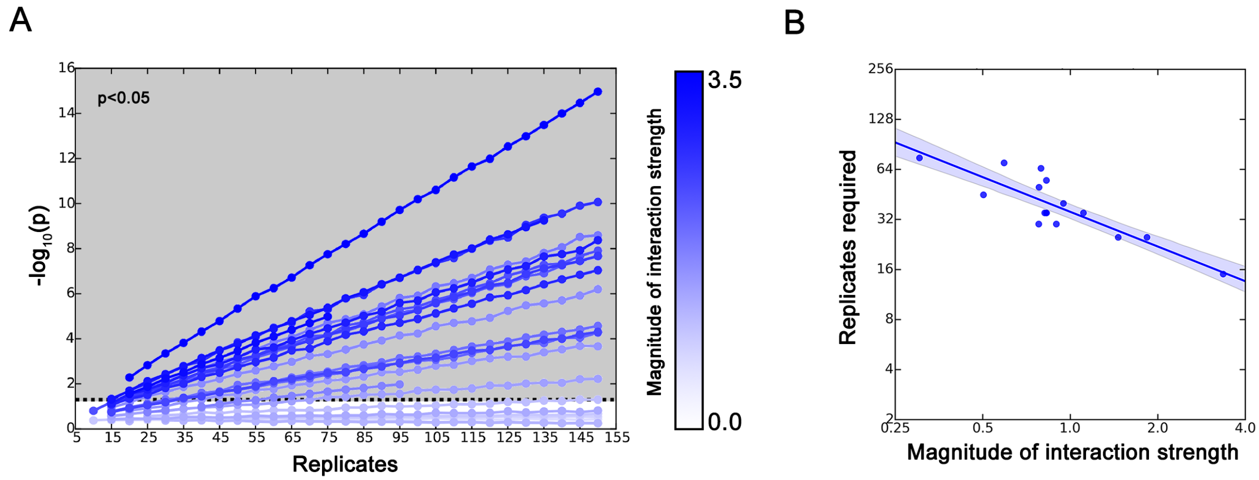


**Fig. S6. Number of droplets (replicates) required to infer microbial interactions of varying strengths using MINI-Drop. (a)** Scatter plot of the number of droplets (replicates) vs. the statistical significance of the interaction ($-\log_{10}(p)$) based on the Mann-Whitney U test. Each data point represents an average p-value computed by sub-sampling the data 500 times across a range of depths. The dashed line represents a p-value equal to 0.05 (statistical significance threshold). The gray box represents the parameter regime for statistical significance ($p < 0.05$). The shade of blue represents the magnitude of the inferred interaction using MINI-Drop. **(b)** The number of droplet replicates required for the p-value to be equal to or less than 0.05.

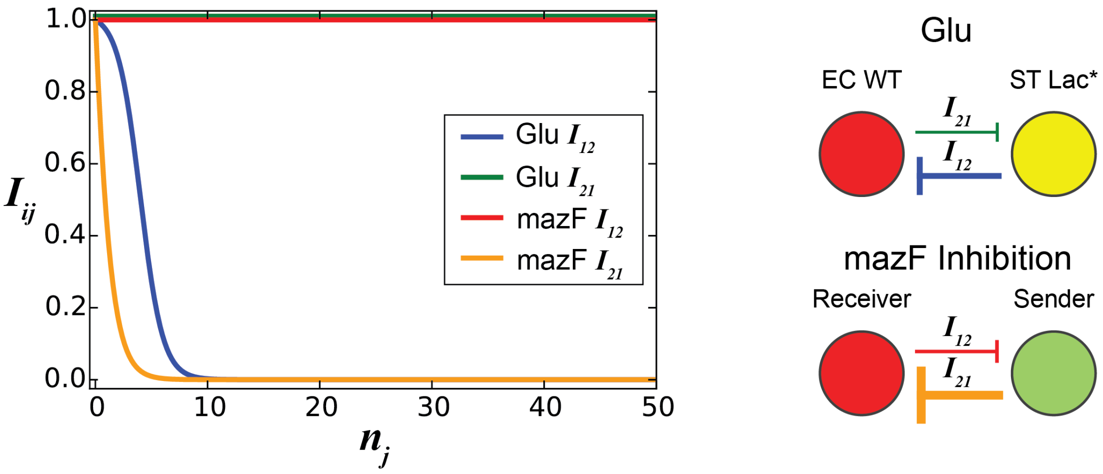


**Fig. S7. Analysis of model interaction strength in two distinct bidirectional negative interaction networks**. Inferred interaction network topologies of pairwise consortia in Experiments E2 and E5 (**Table 1**) were modeled using the discrete-time Markov growth model. The inter-strain interactions in the model are represented as $I_{ij}$ (see Materials and Methods). The toxin mediated negative interaction (orange, right) exhibited a higher sensitivity to partner cell number in the model compared to the ST Lac* to EC WT negative interaction (blue, right).
